# AlphaPept, a modern and open framework for MS-based proteomics

**DOI:** 10.1101/2021.07.23.453379

**Authors:** Maximilian T. Strauss, Isabell Bludau, Wen-Feng Zeng, Eugenia Voytik, Constantin Ammar, Julia Schessner, Rajesh Ilango, Michelle Gill, Florian Meier, Sander Willems, Matthias Mann

**Affiliations:** Department of Proteomics and Signal Transduction, Max Planck Institute of Biochemistry, Martinsried, Germany; NNF Center for Protein Research, Faculty of Health Sciences, University of Copenhagen, Copenhagen, Denmark; Nvidia Corporation, Santa Clara, CA, USA; OmicEra Diagnostics GmbH, Planegg, Germany; Functional Proteomics, Jena University Hospital, Jena, Germany

**Keywords:** Mass spectrometry, Python, open-source, proteomics, open-source, search algorithm, proteome informatics

## Abstract

In common with other omics technologies, mass spectrometry (MS)-based proteomics produces ever-increasing amounts of raw data, making their efficient analysis a principal challenge. There is a plethora of different computational tools that process the raw MS data and derive peptide and protein identification and quantification. During the last decade, there has been dramatic progress in computer science and software engineering, including collaboration tools that have transformed research and industry. To leverage these advances, we developed AlphaPept, a Python-based open-source framework for efficient processing of large high-resolution MS data sets. Using Numba for just-in-time machine code compilation on CPU and GPU, we achieve hundred-fold speed improvements while maintaining clear syntax and rapid development speed. AlphaPept uses the Python scientific stack of highly optimized packages, reducing the code base to domain-specific tasks while providing access to the latest advances in machine learning. We provide an easy on-ramp for community validation and contributions through the concept of literate programming, implemented in Jupyter Notebooks of the different modules. A framework for continuous integration, testing, and benchmarking enforces solid software engineering principles. Large datasets can rapidly be processed as shown by the analysis of hundreds of cellular proteomes in minutes per file, many-fold faster than the data acquisiton. The AlphaPept framework can be used to build automated processing pipelines using efficient HDF5 based file formats, web-serving functionality and compatibility with downstream analysis tools. Easy access for end-users is provided by one-click installation of the graphical user interface, for advanced users via a modular Python library, and for developers via a fully open GitHub repository.

## INTRODUCTION

Increasingly large data sets, combined with exponentially increasing computational power and algorithmic advances, are transforming every aspect of science. This is accompanied and enabled by developments in open and transparent science. The open-source community has been a particular success, starting as a fringe movement to a recognized standard for software development, whose value is embraced and adapted even by the largest technology companies. Public exposure supports high code quality through scrutiny by developers from diverse backgrounds, while increasingly sophisticated collaboration mechanisms allow rapid and robust development cycles. The most advanced machine and deep learning research, for example, builds on open-source projects and datasets and is itself open-source. These laudable developments reflect the core ideas of science and present great opportunities in the ever more important computational fields.

In mass spectrometry (MS)-based proteomics, algorithms and computational frameworks have been a cornerstone in interpreting the data, resulting in a large variety of different proteomic software packages and algorithms, ranging from commercial, freely available to open source, exemplified by and reviewed in (Välikangas, Suomi, and Elo 2017; Chen et al. 2020). Typical computational workflows comprise the detection of chromatographic features, peptide spectrum matching, all the way through protein inference and quantification (Nesvizhskii, Vitek, and Aebersold 2007; Zhang et al. 2020). Advances in (MS)-based proteomics are also being accelerated through the sharing of datasets, such as publicly available data on the Proteome Exchange repository (Vizcaíno et al. 2014; Deutsch et al. 2017).

Prompted by the developments in the Python scientific environment and in collaborative development tools, we developed AlphaPept, a Python-based open-source framework for efficient processing of large amounts of high-resolution MS data. Our main design goals were accessibility, analysis speed, and robustness of the code and the results. Accessibility refers to the idea of facilitating the contribution of algorithmic ideas for (MS)-based proteomics, which is today typically limited to bioinformatics experts. We decided on Python because its clear, easy-to-understand syntax, and because the excellent supporting scientific libraries make it easier for developers from different backgrounds to contribute to and implement new ideas. Using community-tested packages makes the codebase more maintainable and robust, allowing us to focus on domain knowledge instead of implementation details. We furthermore adopted a recent implementation of ‘literate programming’ (Knuth 1984), in which code and documentation are intertwined. Using the nbdev package, the codebase is connected to extensive documentation in Jupyter Notebooks in a way that immediately explains the algorithmic background, making it easier to understand the underlying principles and documenting design decisions for others (Kluyver et al. 2016). With the help of the Numba package for just-in-time compilation (JIT) of Python code (Lam, Pitrou, and Seibert 2015), AlphaPept achieves extremely fast computation times. Furthermore, we implemented robust design principles of software engineering on GitHub, such as continuous integration, deployment and extensive automated validation.

Depending on the user, AlphaPept can be employed in multiple ways. A ‘one-click’ installer can be freely downloaded for Windows, providing a web server-based graphical user interface (GUI) and a command line interface; A Python library that allows re-use and modification of its functionality in custom code, including in Jupyter Notebooks that have become a standard in data science and finally, in a scalable could environment.

In the remainder of the paper, we describe the functionality of AlphaPept on the basis of nbdev notebooks, such as feature finding, peptide identification and protein quantification. We demonstrate the capabilities of AlphaPept on small- and large-scale datasets. Finally, we demonstrate how AlphaPept can be utilized as a proteomic workflow management system and how it can be integrated with downstream analysis tools such as Perseus or the Clinical Knowledge Graph (CKG), (Santos et al. 2020; Tyanova et al. 2016) and we provide an outlook on novel functionality to be incorporated soon.

## RESULTS

### Overview of AlphaPept architecture

Academic software development is often highly innovative but is rarely undertaken with dedicated funding or long term personnel stability. Such constraints have successfully been mitigated by collaborative software engineering approaches and the collective efforts of volunteers. This is exemplified in state of the art open-source projects such as NumPy (Harris et al. 2020) and scikit-learn (Pedregosa et al. 2011). This paradigm has also been taken over by relatively recent and highly popular deep learning frameworks like Google’s Tensorflow (Martín Abadi et al. 2015) and Facebook’s PyTorch (Paszke et al. 2019) and is thought to lead to increased code quality due to community exposure and a large testing audience. Inspired by these developments, AlphaPept implements robust design principles of software engineering on GitHub, such as continuous testing and integration. For instance, code contributions can be made via pull requests which are automatically validated. By making the code publicly available and providing a stringent testing environment, we hope to encourage contribution and testing from a diverse background while maintaining very high code quality.

Organization in notebooks with nbdev allows us to collect documentation, code and tests in one place. This enables us to automatically generate the documentation, extract production code and test functionality by executing the notebooks. Furthermore, we extend the notion of unit and system testing by including real world data sets on which the overall improvement of newly implemented functionality is routinely evaluated. To continuously monitor system performance, summary statistics are automatically uploaded to a database where they are visualized in a dashboard.

The advantages of high-level languages generally come at the price of execution speed, especially for Python. As a result, this expressive language is often only used as a thin wrapper on C++ libraries. In AlphaPept, we make use of the Numba project (Lam, Pitrou, and Seibert 2015), which allows us to compile our Python algorithms directly with the industry-standard LLVM compiler (backend to most C++ compilers and supercomputing languages such as Julia). This allows us to speed up our code by orders of magnitude without losing the benefits of the intuitive Python syntax. Furthermore, AlphaPept readily parallelizes computationally intensive parts of the underlying algorithms on multiple CPU cores or – if available – Graphical Processor Units (GPUs) for further performance gains.

As far as possible, AlphaPept uses the standard, but powerful packages of the Python data analysis universe, namely NumPy for numerical calculations, pandas for spreadsheet-like data structures and scikit-learn for machine learning (Fig. 1A). Furthermore, we chose the binary, high performance HDF5 file format, which is used across scientific areas, including ‘big data’ projects (see below). All these packages are platform-independent, allowing deployment of AlphaPept on Windows, Mac and Linux computers, including cloud environments.

**Figure 1:**
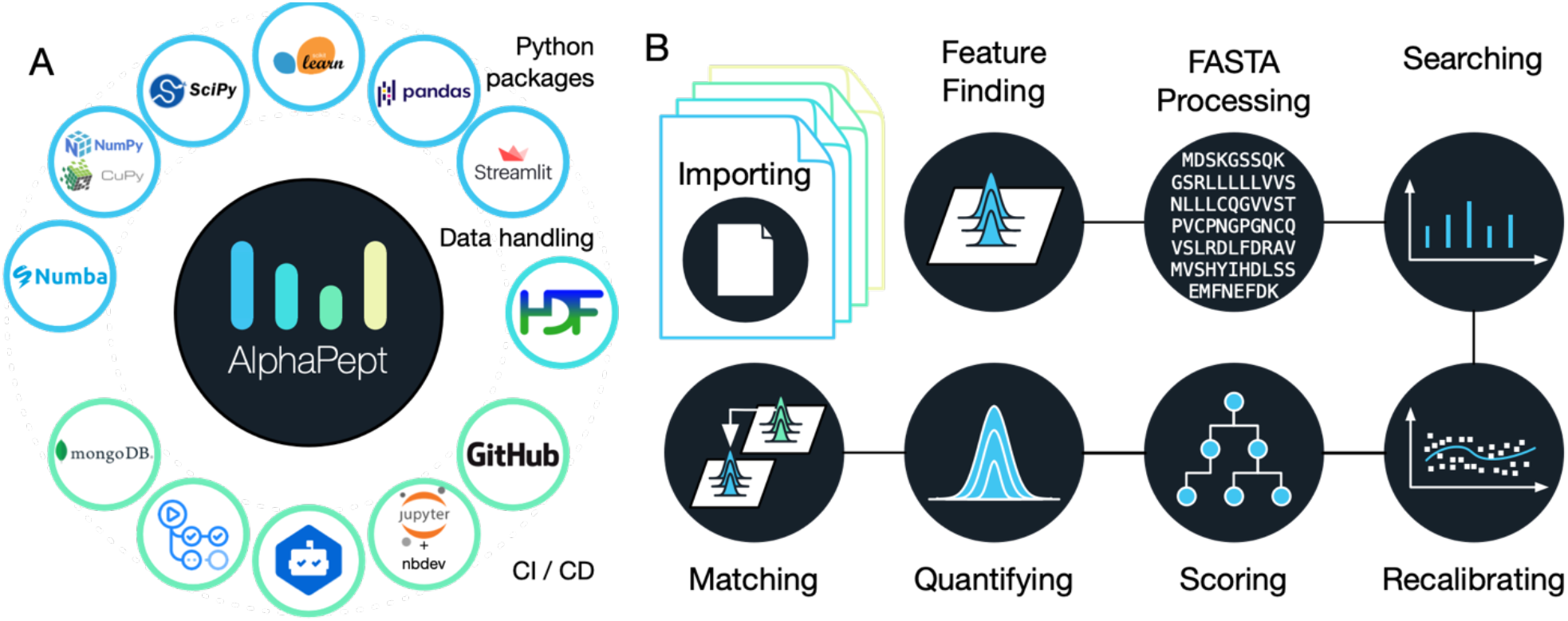
AlphaPept ‘ecosystem’ and Modules. **A** AlphaPept relies on multiple community-tested packages. We use highly optimized libraries such as Numba, NumPy, CuPy, scikit-learn, SciPy and pandas to achieve performant code. As GUI, we provide a browser-based application built on streamlit. For data handling, the HDF5 file technology is used. The repository itself is hosted on GitHub, the core code is documented in Jupyter Notebooks using the nbdev package. To ensure maintainability, packages are continuously monitored for updates via dependabot. New code is automatically validated using GitHub actions and summary statistics (timing, identifications and quantifications) are uploaded to a mongoDB database and visualized. **B** All algorithmic code of AlphaPept is organized in Jupyter Notebooks. For the key processing steps in the pipeline, such as importing raw data, Feature Finding, FASTA processing, Searching, Recalibrating, Scoring, Quantifying and Matching, there are individual notebooks with background information and the code.

An integral feature of AlphaPept development are Jupyter notebooks, which have become ubiquitous in scientific computing. Using the nbdev package, each part of the MS-based proteomics workflow is modularized into a separate notebook. This allows extensive documentation of the underlying algorithmic production code, which is automatically extracted from and synchronized with the notebooks. Furthermore, the notebooks capture the background information of each part of the computational proteomics workflow, making it much easier to understand the underlying principles. We have found this to be an excellent way of developing software, which brings together the typical cycle of exploration in notebooks with the production of a robust and tested code base. Figure 1B shows an overview of the steps in the analysis of a typical proteomics experiment in AlphaPept corresponding to the notebooks. These separate processing steps will be discussed in turn in the sections below.

### Highly efficient and platform-independent MS data access

MS-based proteomics or metabolomics generates complex data types of MS1 level features, variable length MS2 data and mappings between them. Furthermore, data production rates are rapidly increasing, making robust and fast access a central requirement. The different MS vendors have their own file formats, which may be highly optimized but are meant to be accessed by their own software. We therefore faced the task of extracting the raw data into an equally efficient but vendor-neutral format that could be accessed rapidly.

First, AlphaPept needs to convert vendor specific raw files. For Thermo files we created a cross-platform Python application programming interface (API) that can directly read .RAW MS data (pyRawFileReader, Fig. 2a). It uses PythonNET for accessing Thermo’s RawFileReader .NET library (Zeng, Wen-Feng 2021, 1), obviating the need for Thermo’s propietory MSFileReader. For Windows, PythonNET is available by default as a part of Windows’ .NET Framework. For Linux and MacOS, PythonNET requires the open-source Mono library. Although our solution uses stacked APIs, loading the spectra of a Thermo .RAW file of 1.6 Gb into RAM takes only about one minute which can be speeded up even more by parallel file processing. Access to Bruker’s timsTOF raw data is also directly handled from our Python code, in this case through a wrapper to the external timsdata.dll C/C++ library, both made available by Bruker. In parallel with this publication, we provide AlphaTims, a highly efficient package to access large ion mobility time-of-flight data through Python slicing syntax and with ultra-fast access times (https://github.com/MannLabs/alphatims).

**Figure 2:**
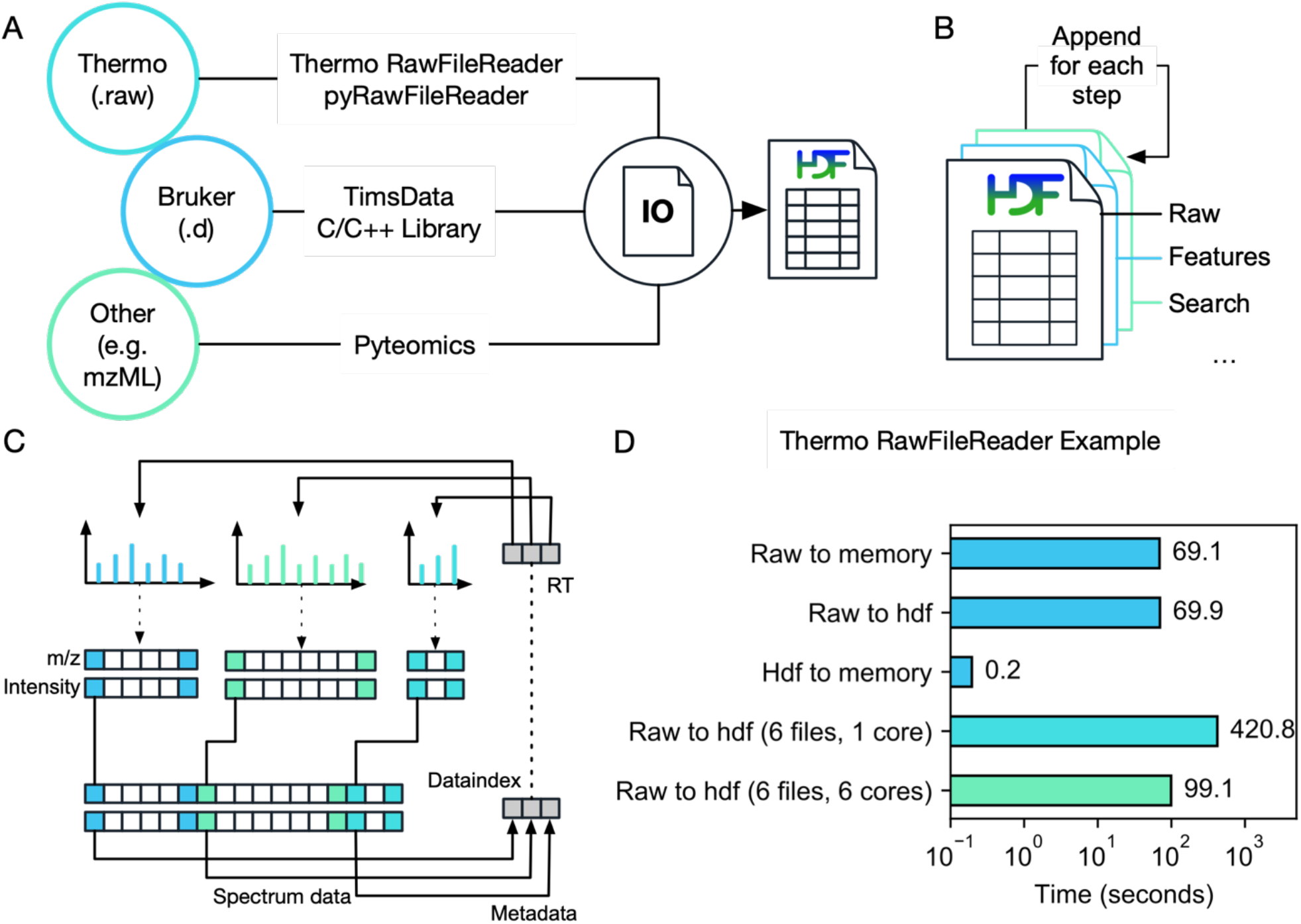
Highly efficient and platform-independent MS data access. **A** MS data from different vendors is imported to an HDF5 container for fast and platform-independent data access. To read Thermo data, we provide a Python application programming interface. Bruker data is accessed via Bruker’s proprietary DLL. Additionally, generic data can be imported using the Pyteomics package. **B** The output of each processing step is appended to the HDF5, allowing processing in a modular way. **C** To efficiently store MS spectra, multiple spectra of variable length are concatenated, and start indices are saved in a lookup table. **D** HDF5 Accessing times. Loading data from HDF5 into memory takes less than 1s for a typical 2h full proteome analysis of a HeLa sample acquired on a Thermo Orbitrap mass spectrometer.

To accommodate raw data acquired through other vendors, we use Pyteomics (Goloborodko et al. 2013; Levitsky et al. 2019). This package allows reading mzML and other standard MS data formats with Python. Thus, by first converting raw data with external software such as e.g. MSConvert (Adusumilli and Mallick 2017), AlphaPept also provides a generic framework for all vendors.

As a storage technology, we chose HDF5 (Hierarchical Data Format 5), a standard originally developed for synchrotron and other extremely large scale experimental data sets, that has now become popular in a wide range of scientific fields (Folk et al. 2011). HDF5 has many benefits such as independence of operating systems, arbitrary file size, extremely fast accession and a transparent, flexible data structure. The latter is achieved by organizing HDF5 files in groups and subgroups, each containing arrays of arbitrary size and metadata which describes these arrays and (sub)groups. In the last few years, it is also becoming more popular in the field of MS (Wilhelm et al. 2012, 5). AlphaPept adopts the HDF5 technology via the Python’s h5py package (Collette 2013).

As an additional design choice we also store intermediate processing results in the HDF5 container, so that individual processing steps can be performed in a modular way and from different computers. This enables researchers to quickly implement and validate new ideas within the downstream processing pipeline. Thus, for each new sample, AlphaPept creates a new .ms_data.hdf file and for each step in the workflow, the file is extended by a new group (Fig. 2b). In this way, the .ms_data.hdf file ensures full portability, transparency and reproducibility while being fast to access and with minimal storage requirements. For example, the 1.6 Gb Thermo file mentioned above is converted to a HDF5 file of 200 MB, all of which can be accessed in a total of 0.2 s (Fig. 2D).

We next provide functionality for MS data pre-processing, such as centroiding and extraction of the n-most abundant fragments, should this not already have happened in the vendor software. MS1 and MS2 scans form the two major subgroups in the HDF5 file. As HDF5 files are not optimized for lists of arrays with variable length, we convert the many individual spectra into a defined number of arrays, each containing a single data type, but concatenating all spectra. These arrays are organized in two sets: Spectrum metadata (spectrum number, precursor m/z, RT, etc), where each array position corresponds to one spectrum; and spectrum data, where each array position corresponds to a single m/z-intensity pair. To unambiguously match the spectrum datapoints to their metadata, an index array is created. It is part of the first set of arrays and contains a pointer to the position of the first data pair for each spectrum within the second set. The position of the last pair does not need to be stored as it is implied by the start position of the next spectrum. Thereby, all m/z values and intensities for each spectrum can easily be extracted with simple base Python slicing, while fixing the number arrays contained in the hdf container. Loading data from HDF5 to RAMtakes less than a second, effectively speeding up data accession more than 300-fold compared to loading the RAW file (Fig. 2d).

### Extracting isotope features

Having stored the MS peaks from all mass spectra in an efficient data structure, we next determine isotope patterns over chromatographic elution profiles. This computationally intensive task is crucial for subsequent peptide identification and quantification. MaxQuant (Cox and Mann 2008) introduced the use of graphs for feature finding, which was then improved upon by the Dinosaur tools (Teleman et al. 2016) and we also decided to follow this elegant approach.

In the first step – called hill building – centroided peaks from adjacent scans are connected. As there are millions of centroids, our first implementations using pure Python took several minutes of computing time. We subsequently refactored the graph problem and parallelized it for CPUs using Numba and CuPy for GPUs, resulting in a 300-fold speed up (about 1s on GPU). Since not every user has access to GPUs, AlphaPept employs dedicated Python ‘decorators’, a metaprogramming technique allowing a part of the program to modify its another part at compile time to transparently switch between parallelized CPU, GPU and pure Python operation.

In more detail, AlphaPept refines hills by first splitting them in case they have local minima indicating two chromatographic elution peaks (Fig. 3B). Additionally, hills are removed whose elution profiles do not conform to minimal criteria, like minimal length and the existence of local minima. To efficiently connect hills, we compute summary statistics such as weighted average m/z value and a bootstrap estimate of its precision. Hills within retention time boundaries are grouped into pre-isotope patterns. To correctly separate co-eluting features, we generate seeds, which we extend in elution time and check for consistency with a given charge state, similarity in elution profile and for conformity with peptide isotope abundance properties via the averagine model (Senko, Beu, and McLaffertycor 1995). This results in a feature (here a possible peptide precursor mass), which is described by a table.

**Figure 3:**
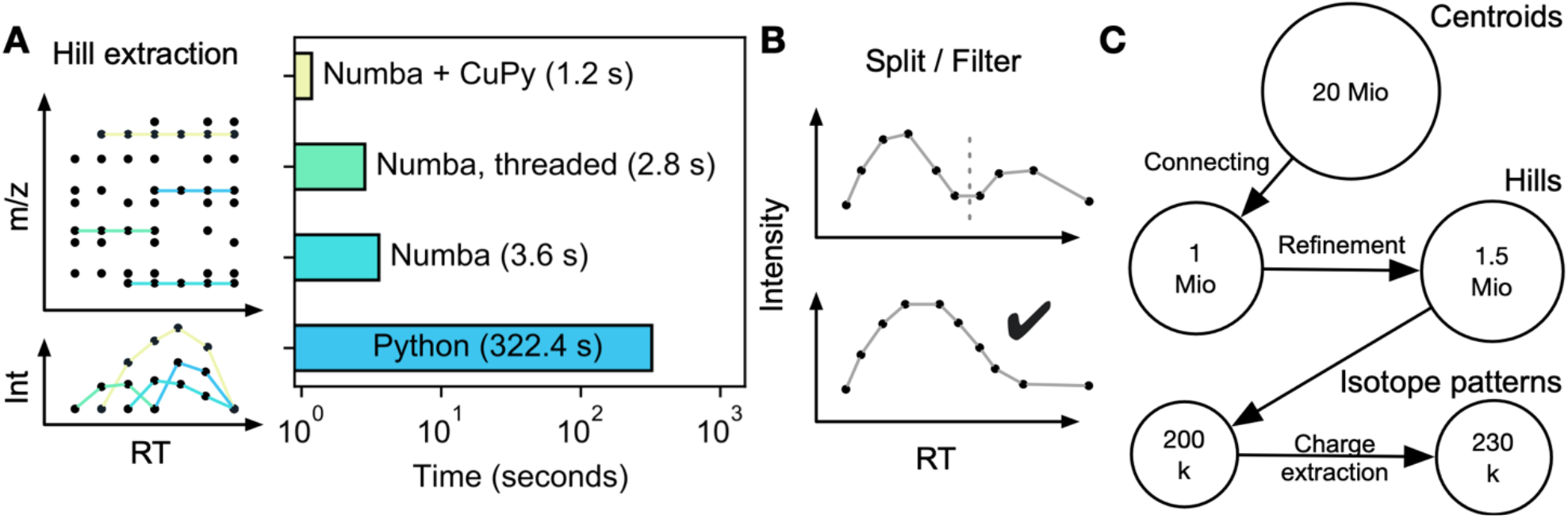
Extracting isotope features. **A** Individual MS peaks of similar masses are connected over the retention time using a graph approach, resulting in ‘hills’. Using a native Python implementation, hill extraction takes several minutes. Numba, parallelization on CPUs or GPUs reduces hill extraction to seconds. **B** Extracted hills are refined by splitting at local minima and only allowing well-formed elution profiles. **C** Starting with 20 million points for a typical Thermo HeLa shotgun proteomics file, these are connected to approximately one million hills, which increased to 1.5 million after hill splitting and filtering. Subsequent processing results in 200,000 pre-isotope patterns that ultimately yield 230,000 isotope patterns due to assignment to specific charge states.

Feature finding on the Bruker timsTOF involves ion mobility as an additional dimension. Currently, this functionality is provided by a Bruker component, which we linked into our workflow via a Python wrapper, and is the only part that is not in natively included as Python code in AlphaPept. Instead, this wrapper uses Python’s subprocess module, which can integrate other tools into AlphaPept just as easily.

For a typical proteomics experiment performed on an Orbitrap instrument, Figure 3C provides an overview of the number of data points from MS peaks to the final list of isotope patterns. Note that AlphaPept can perform feature finding separately for each file as soon as it is acquired (described below). Furthermore, although described here for MS1 precursors, the AlphaPept feature finder is equally suited to MS2 data that occur in parallel reaction monitoring (PRM) or DIA acquisition modes.

### Peptide spectrum matching

The heart of a proteomics search engine is the matching of msms spectra to peptides in a protein sequence database. AlphaPept parses FASTA files containing protein sequences and descriptions, ‘digests’ them into peptides and calculates fragment masses according to user specified rules and amino acid modifications (Fig 3D). We again use HDF5 files, which enables efficient storage of fragment series despite their varying lengths. Generation of this database only happens once per project and only takes minutes for typical organisms and modifications. From a FASTA file of the human proteome, typically five million ‘in silico’ spectra of fragment masses are generated. In case no enzyme cleavage rules are specified or for open search with wide precursor mass tolerances, the fragments are instead generated on the fly to avoid excessive file sizes.

To achieve maximum speed, AlphaPept employs a very rapid fragment counting step to determine initial peptide spectrum matches (PSMs). As this step only involves addition and subtraction of elements in numerical arrays, the machine code produced by Numba is very efficient and easily parallelized. This leaves a much smaller number of peptides that have at least a minimum number of fragment matches to the experimental spectrum. (This is similar to the Morpheus score (Wenger and Coon 2013), which also computes the fraction of msms signals accounted for by the match.) For the human proteome and mass measurement accuracy of parts per million, the initial millions of comparisons are decreased to a maximum of top-n remaining candidates per msms spectrum (typically 10). This enables more computationally expensive scoring in a second step. Different scores can be implemented in AlphaPept, and by default we chose the widely used X!Tandem score (Craig and Beavis 2003). Note that the sole function of this score is to rank the PSMs, whereas statistical significance is determined by counting reverse database hits and by machine learning (see below).

We perform a first search for the purpose of recalibrating the mass scale as a function of elution time (Fig. 4B). Here, we use weighted nearest neighbor regression instead of binning by retention time (explained in the accompanying Jupyter Notebook). The k-nearest neighbors regressor that we selected allows non-linear grouping in several dimensions simultaneously (retention time and mass scale in the case of Orbitrap data and additionally ion mobility in the case of timsTOF data).

**Figure 4.**
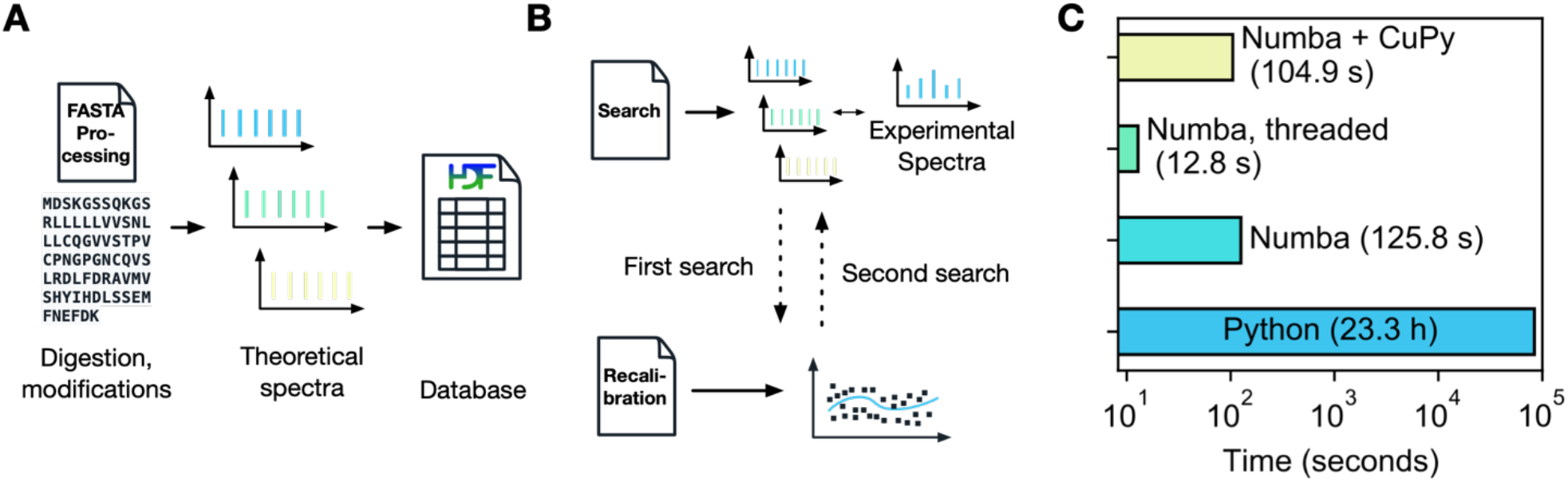
Database search. **A** The FASTA processing notebook contains functionality to calculate fragment masses from FASTA files which are saved in an HDF5 container for subsequent searches. **B** Initially, a first search is performed, and masses are subsequently recalibrated. Based on this recalibration, a second search with more stringent boundaries is performed. **C** Using the decorator strategy, the search can be drastically speeded up, from 23 h in a pure Python implementation to seconds with Numba and CuPy.

**Figure 5:**
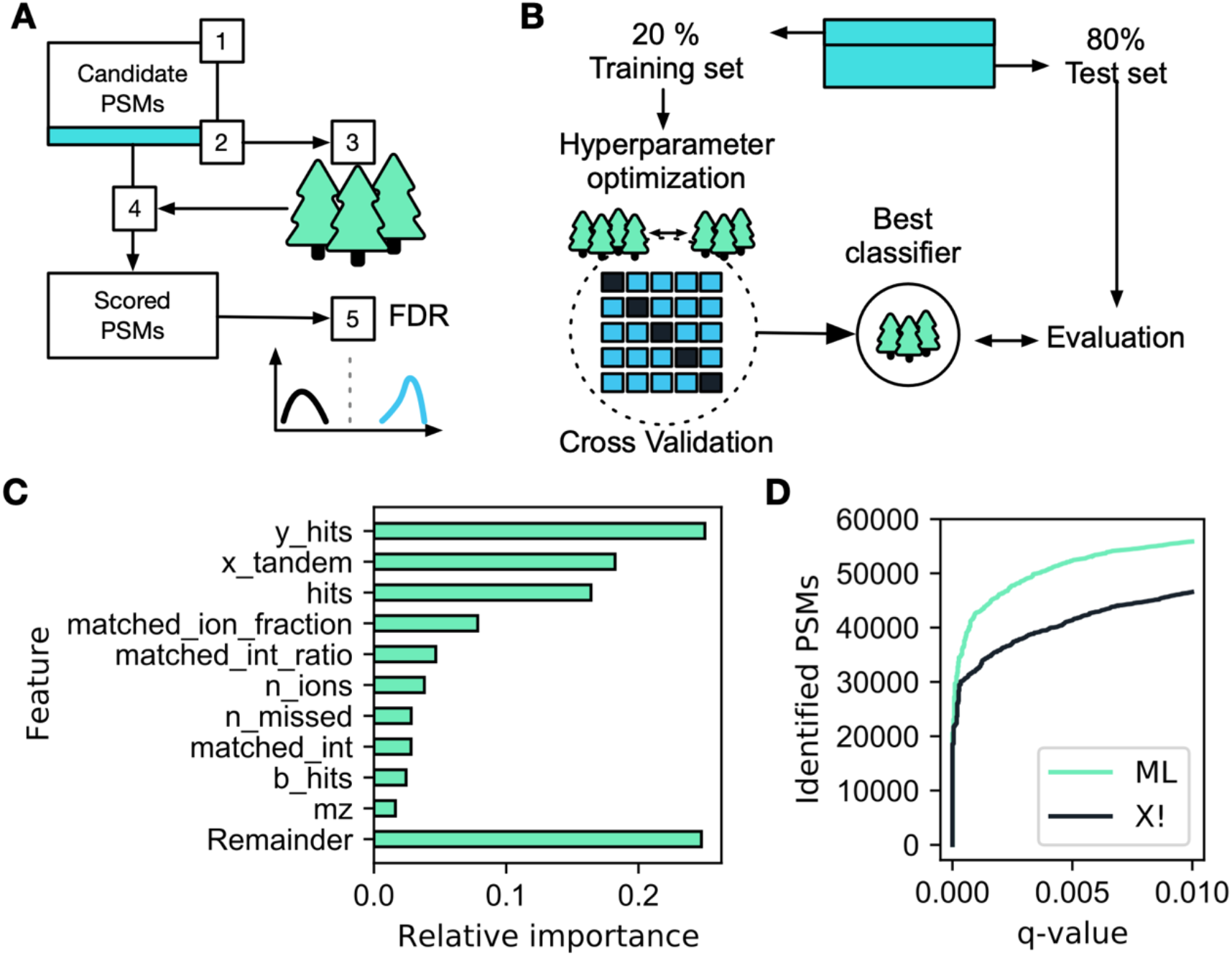
Machine learning-based scoring and FDR estimation. **A** We train a Random Forest (RF) classifier on a subset of candidate PSMs to distinguish targets from decoys based on PSMs characteristics. A semi-supervised machine learning model is applied with the following steps: (1) extraction of all candidate PSM scores, (2) selection of a PSM subset for machine learning, (3) training of a RF classifier, and (4) application of the trained classifier to the full set of PSM candidates. Finally, the probability of the RF prediction is used as a score for subsequent FDR control (5). **B** Training of the classifier (step 4 in panel A) follows a train-test split scheme where only a fraction of the candidate subset is used for training. Using stringent cross-validation, multiple hyperparameters are tested to achieve optimal RF performance. The best classifier is benchmarked against the remaining test set. **C** Example feature importance for an Orbitrap test set, where the number of y-ion hits is the highest contributing factor to the model. Note that the RF algorithm can utilize any database identification score such as the X!Tandem score chosen here, which is the second most important feature. See the *AlphaPept workflow and files* Notebook for an explanation of features. **D** Optimized identification with the ML score. Compared to the X!Tandem score alone, the ML optimization identified about 15% more PSMs for the same q-value.

Having recalibrated the data, the main search is performed with an adapted precursor tolerance. We furthermore calculate the matched ion intensity, matched ions, neutral loss matches for further use and reporting together with charge, retention time and other data.

To demonstrate the speed up achieved by our architecture and the performance decorator, we timed illustrative examples (Fig. 4C). On a HeLa cell line proteome acquired in a single run, comparing 260k spectra to 5 million database entries, the computing time in pure Python was about 23 h. This decreased to 126 s when employing Numba (> 500x improvement), to 105 s when using Numba with CuPy on GPU and further to 13 s on multi-threaded CPU (see companion Figure Notebook). The GPU acceleration is not larger because the code is already very efficient on CPU and some workflow tasks are memory bound instead of computationally bound. Improved memory management on GPU could further decrease GPU computational time. In any case, AlphaPept reduces the PSM matching step to an insignificant part total computation time.

### Machine learning based scoring and FDR estimation

Assessing the confidence of PSMs requires a scoring metric that separates true (correctly identified) from false (wrongly identified) targets in the database. Multiple defined features are calculated by the AlphaPept search engine and used in a score to rank the targets. A nonsense database of pseudo-reversed sequences where the terminal amino acid remains unchanged (de Godoy et al. 2008) is used to directly estimate the False Discovery Rate (FDR) by counting reverse hits. Score thresholds subsequently decide which targets should be considered identified. To further validate this approach and to ensure accurate FDR estimation across different development stages in AlphaPept, our GitHub testing routine includes an empirical two species FDR test based on an ‘entrapment strategy’ (Muntel et al. 2019).

In recent years, machine learning has gained increasing momentum in science in general, but also in its specific applications to MS data analysis. One of the first of these was the combination of multiple scoring metrics to a combined discriminant score that best separates high scoring targets from decoys. This was initially integrated into PSM scoring through an external reference dataset to train the classifier (Keller et al. 2002). The widely used Percolator approach subsequently employed a semi-supervised learning approach that was trained directly on the dataset itself (Käll et al. 2007). This automatically adapts the ML model to the experimental data and along with other MS analysis tools (MacLean et al. 2010; Röst et al. 2014; Teleman et al. 2015; Fondrie and Noble 2021; Rosenberger et al. 2017) we also employ semi-supervised learning for PSM scoring in AlphaPept.

The AlphaPept scoring module falls into five parts: (1) feature extraction for all candidate PSMs, (2) selection of a candidate subset, (3) training of a machine learning classifier, (4) scoring of all candidate PSMs and (5) FDR estimation by a target-decoy approach (Fig. 4A). Most features for scoring the candidate PSMs are directly extracted from the search results, such as the number of b- and y-ion hits and the matched ion intensity fraction. Some additional features are subsequently determined, including the sequence length and the number of missed cleavages. After feature extraction, a subset of candidate PSMs is selected with an initial 1% FDR threshold based only on the X!Tandem score (Fig. 4B). Together with an equal number of randomly selected decoys, this creates a balanced dataset for machine learning. This is split into training and test sets (20% vs. 80%) and provides the input of a ML classifier. We chose a standard scikit-learn random forest classifier as it performed similarly to XGBoost with fewer dependencies on other packages. We first identify optimal hyper-parameters for the classifier with a grid-search via five-fold cross-validation. The resulting best classifier optimally separates target from decoy PSMs on the test set. Applying the trained classifier to the entire set of candidate PSMs yields discriminant scores that are used to estimate q-values based on the classical target-decoy competition approach.

The contribution of different features to the discriminant score for an exemplary tryptic HeLa sample is shown in Figure 4C. Interestingly, for our data, the number of matched y-ions alone outperforms the basic search engine score and most of the top-ranking features are related to the number of matched ions and their intensity. The ML algorithm markedly improved the separation of targets vs decoys, retrieving a larger number of PSMs at every q-value (Fig. 4D). ML-based scoring in AlphaPept improved identification rates by 15% at a 1% FDR at the PSMs level, in line with previous efforts (Käll et al. 2007). AlphaPept allows ready substitution of the underlying PSM score and machine learning algorithms. Furthermore, additional features to describe the PSMs are readily integrated, such as ion mobility or predicted fragment intensities. We envision that this kind of flexibility will enable continuous integration of improved workflows as well as novel ML techniques into AlphaPept.

Once a set of PSMs at a defined FDR is identified, protein groups are determined via the razor protein approach (Nesvizhskii and Aebersold 2005). Here, peptides that could potentially map to multiple unique proteins are assigned to the protein group that already has most peptide evidence. We determine protein-level q-values by selecting the best scoring precursor per protein, followed by FDR estimation by target-decoy competition similar to the peptide level (Nesvizhskii 2010; Savitski et al. 2015; The et al. 2016; Gupta and Pevzner 2009). Finally, we validated the scoring and FDR estimation in AlphaPept with the entrapment strategy mentioned above, by analyzing a HeLa sample with a mixed species library, containing targets and decoys derived from both a human FASTA and a FASTA from Arabidopsis thaliana. This revealed that AlphaPept provides accurate q-value estimates, reporting approximately the same number of *Arabidopsis thaliana* proteins as decoy proteins at 1% protein FDR.

### Label-free quantification

The ultimate goal of a proteomics experiment is to derive functional insights or assess biomarkers from quantitative changes at the protein level, to which peptide identifications are only means to an end. Algorithmically this quantification step entails either the determination of isotope ratios in the same scans (for instance SILAC, TMT or EASI-tag ratios) or the somewhat more challenging problem of first integrating peaks and then deriving quantitative ratios across samples (label-free quantification), which we focus on here. We initially adapted the MaxLFQ pipeline for label-free quantitative proteomics data (Cox et al. 2014). The first task is to determine normalization factors for each run as different LC MS/MS runs need to be compared – potentially spaced over many months in which instrument performance may vary – and as total loading amounts likewise vary for instance due to pipetting errors. The basic assumption is that the majority of peptides are not differentially abundant between different samples. This allows deriving the run-specific normalization factors by minimizing the between-sample log peptide ratios (Cox et al. 2014) (Note that this assumption is not always valid and can be restricted to certain protein classes.). In a second step, adjusted intensities are derived for each protein, such that protein intensities between different MS runs can be compared. To this end we derive the median peptide fold changes that maximize consistency with the peptide evidence.

The normalization, as well as protein intensity profile construction, are quadratic minimization problems of the normalization factors or the intensities, respectively. Such minimization problems can be solved in various ways but one fundamental challenge is that these algorithms have a time complexity of O(n^2^), meaning that the computation time increases quadratically with the number of comparisons. One strategy to overcome this limitation is to only perform minimization on a subset of all possible pairs (termed ‘FastLFQ’) (Cox et al. 2014). Despite this, the computation time of the underlying solver will determine the overall runtime and accounts for the long run times on very large datasets. However, a variety of very efficient solvers that are based on different algorithms are contained in the Python SciPy package (SciPy 1.0 Contributors et al. 2020). To test these approaches, we created an *in silico* test dataset with a known ground truth (see Quantification Notebook). Comparing different solvers using our benchmarking set uncovered dramatic differences in precision, runtime and success rate (Fig. 6A). Among the better performing algorithms were the least-squares solvers that were previously used. The *Broyden–Fletcher–Goldfarb–Shanno* (L-BFGS-B), *Sequential Least Squares Programming* (SLSQP) and *Powell* algorithms were particularly fast and robust solutions being up to 16x quicker than the Trust Region Reflective algorithm (trf) from the default least-squares solver. More remarkably, they were able to optimize much better to our known ground truth. Of all four tested optimizers, the mean error of trf was, on average 24% worse. Being able to readily switch between different solvers provided by SciPy allows us to fall back on other solvers if the default solver fails, i.e. AlphaPept will switch from L-BFGS-B to Powell if the solution does not converge.

**Figure 6:**
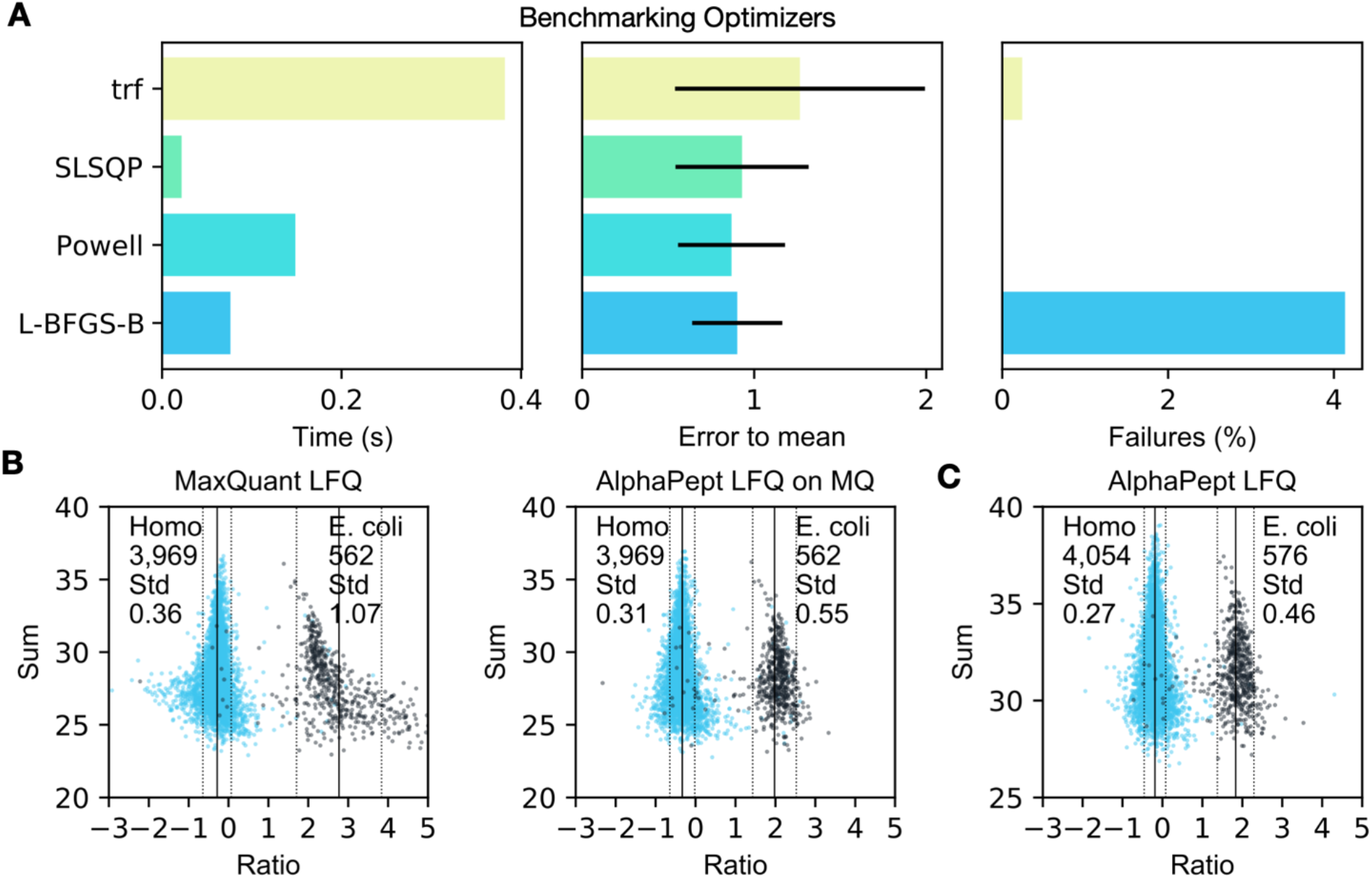
Algorithm selection and performance of label-free quantification. **A** Timings of different, highly optimized solvers from the SciPy ecosystem, to extract optimal protein intensity ratios in AlphaPept. Solvers showed drastic differences in speed, closeness to ‘ground truth’, and proportion of successful optimizations on *in-silico* test data. Based on these tests, AlphaPept employs a hybrid optimization strategy that uses L-BFGS-B and Powell for optimized performance, robustness and speed. **B** Comparing the AlphaPept LFQ solver on MaxQuant output data demonstrates similar separation in mixed-species datasets with smaller standard deviations. **C** Applying AlphaPept directly on the same dataset further improves identifications and quantification accuracy.

**Figure 7:**
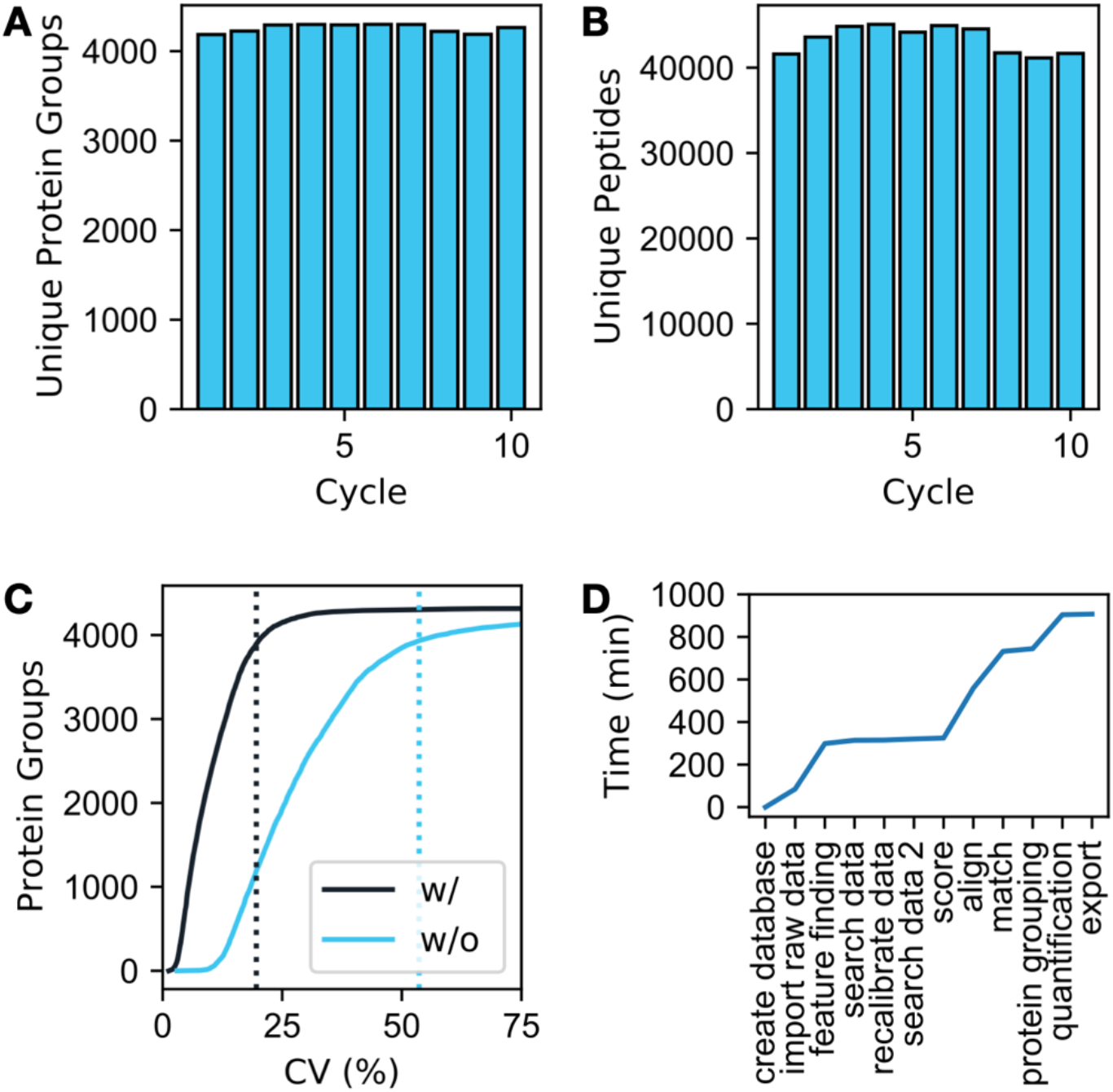
Benchmarking AlphaPept on 200 HeLa proteomes. A total of 200 DDA HeLa cell proteomes – the 10 cycle long term performance test from Kuster and coworkers (181 Gbyte) (Bian et al. 2020)– was analyzed by AlphaPept. **A** Identification performance at the protein group level. **B** Identification performance at the peptide level. **C** Quantification performance with or without MaxLFQ optimization. For 90% of protein groups, CVs are below 20% and 54%, respectively. **D** Timing of the AlphaPept computational pipeline. Search through scoring are highly optimized and contribute little to overall computation time.

We compared our method to MaxLFQ in a quantitative two-species benchmarking dataset, in which *E. coli* proteins change their abundance by a factor of six between conditions, while human proteins do not change (Meier et al. 2018). To specifically assess the benefits of the new optimization strategy, we first tested the algorithm directly on the MaxQuant output (see companion Notebook for Figure 6). Both approaches clearly separated human and *E. coli* proteins, however, the standard deviation was smaller when applying the AlphaPept optimization algorithm, which also has fewer outlier quantifications (Fig 6B), supporting the analysis of the *in-silico* test set. Comparing results of the complete workflow with AlphaPept on the same files further improved identifications and quantifications.

### Match-between-runs (MBR) and dataset alignment

We implemented functionality to transfer the identifications of MS1 features to unidentified MS1 features of other runs (*match-between-runs*). First, we align multiple datasets on top of each other by applying a global offset in retention time, mass and – where applicable – ion mobility. To determine offsets for all runs, we first compare all possible pairs of runs and calculate the median offset from one dataset to another based on the precursors that were identified in both. As these offsets are linear combinations of each other, i.e., the offset from dataset A to dataset C should be the offset from dataset A to B and B to C; this becomes an overdetermined equation system, which we solve by a weighted linear regression model with the number of shared precursors as weights.

After dataset alignment, we group precursors of multiple runs and determine their expected properties as well as their probability density and create a library of precursors. Next, we take the unidentified MS1 features from each run and extract the closest match from the library of precursors. Finally, as we know the probability density of each feature, we can calculate the Mahalanobis distance from each identification transfer and use this as a probability estimate to assess the likelihood that a match is correct. Further information about the alignment and matching algorithm can be found in the Matching notebook.

### Benchmarking AlphaPept on large data sets

A prime goal of the AlphaPept effort is robustness and speed. To showcase the usability of AlphaPept for large scale studies we re-analyzed 200 HeLa proteomes from a recently published long-term performance test (Bian et al. 2020). To confirm comparable identification performance in the initial analysis, which was done with MaxQuant, we evaluated the number of uniquely identified protein groups and PSMs per group. This yielded a median of 4277 unique protein groups and 43,872 unique peptides per experimentally defined group, as expected. Next, we compared the protein level quantification. The median coefficient of variation without our Python maxLFQ implementation was 27.1% and 9.2% after LFQ optimization. For 90% of protein groups, CVs were below 20% with LFQ optimization and below 54% without. Investigation of each computational task revealed that a large part is spent on importing raw data and feature finding. Searching and scoring are highly optimized and contribute only a small fraction of the overall computing time. Operations across files such as LFQ alignment and matching again make up a large part of computation time.

### Continuous validation on standard datasets

Our current continuous integration pipeline uses a range of data sets typical for MS workflows. These include standard single shot runs, such as HeLa quality control (QC) runs, as well as recently published studies. For every addition to the main branch of the code base, AlphaPept reanalyzes these files fully automatically, allowing extensive systems checks. Additionally, these checks can be manually triggered at any time and therefore enable swift validation of proposed code changes prior to submitting pull-requests. This makes comparing studies that were analyzed with different software versions much more transparent. To further increase this idea of transparent performance tracking, we automatically upload summary statistics, such as runtime, number of proteins and number of features for each run to a database and visualize these metrics in a dashboard (Extended methods). Table 1 shows example tracking metrics from the database.

**Table 1:**
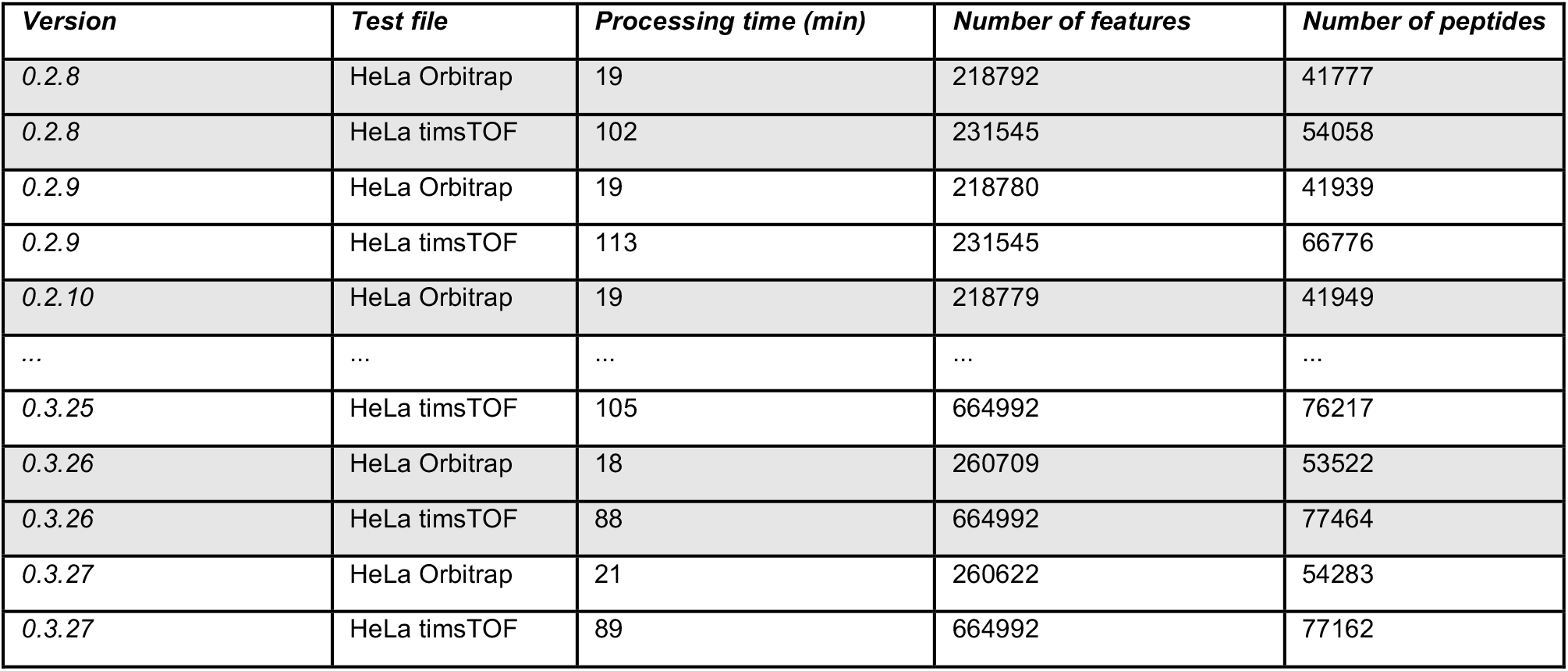
Example performance tracking metrics for different AlphaPept versions extracted from the database.

| **Version** | **Test file** | **Processing time (min)** | **Number of features** | **Number of peptides** |
| --- | --- | --- | --- | --- |
| 0.2.8 | HeLa Orbitrap | 19 | 218792 | 41777 |
| 0.2.8 | HeLa timsTOF | 102 | 231545 | 54058 |
| 0.2.9 | HeLa Orbitrap | 19 | 218780 | 41939 |
| 0.2.9 | HeLa timsTOF | 113 | 231545 | 66776 |
| 0.2.10 | HeLa Orbitrap | 19 | 218779 | 41949 |
| ... | ... | ... | ... | ... |
| 0.3.25 | HeLa timsTOF | 105 | 664992 | 76217 |
| 0.3.26 | HeLa Orbitrap | 18 | 260709 | 53522 |
| 0.3.26 | HeLa timsTOF | 88 | 664992 | 77464 |
| 0.3.27 | HeLa Orbitrap | 21 | 260622 | 54283 |
| 0.3.27 | HeLa timsTOF | 89 | 664992 | 77162 |

### AlphaPept user interface and server

A central element for any software tool is ease of use for the end user. In the most basic setup, this is determined by the accessibility of the GUI. Following recent trends, we decided on server-based technology for AlphaPept. In a basic setup, the web interface is called by connecting to a local server instance on the user’s laptop or local workstation (Fig. 8A) via a browser. For more demanding pipelines, AlphaPept can be run on a powerful processing PC and be accessed from multiple other devices. This makes access to AlphaPept platform independent, including mobile devices.

**Figure 8:**
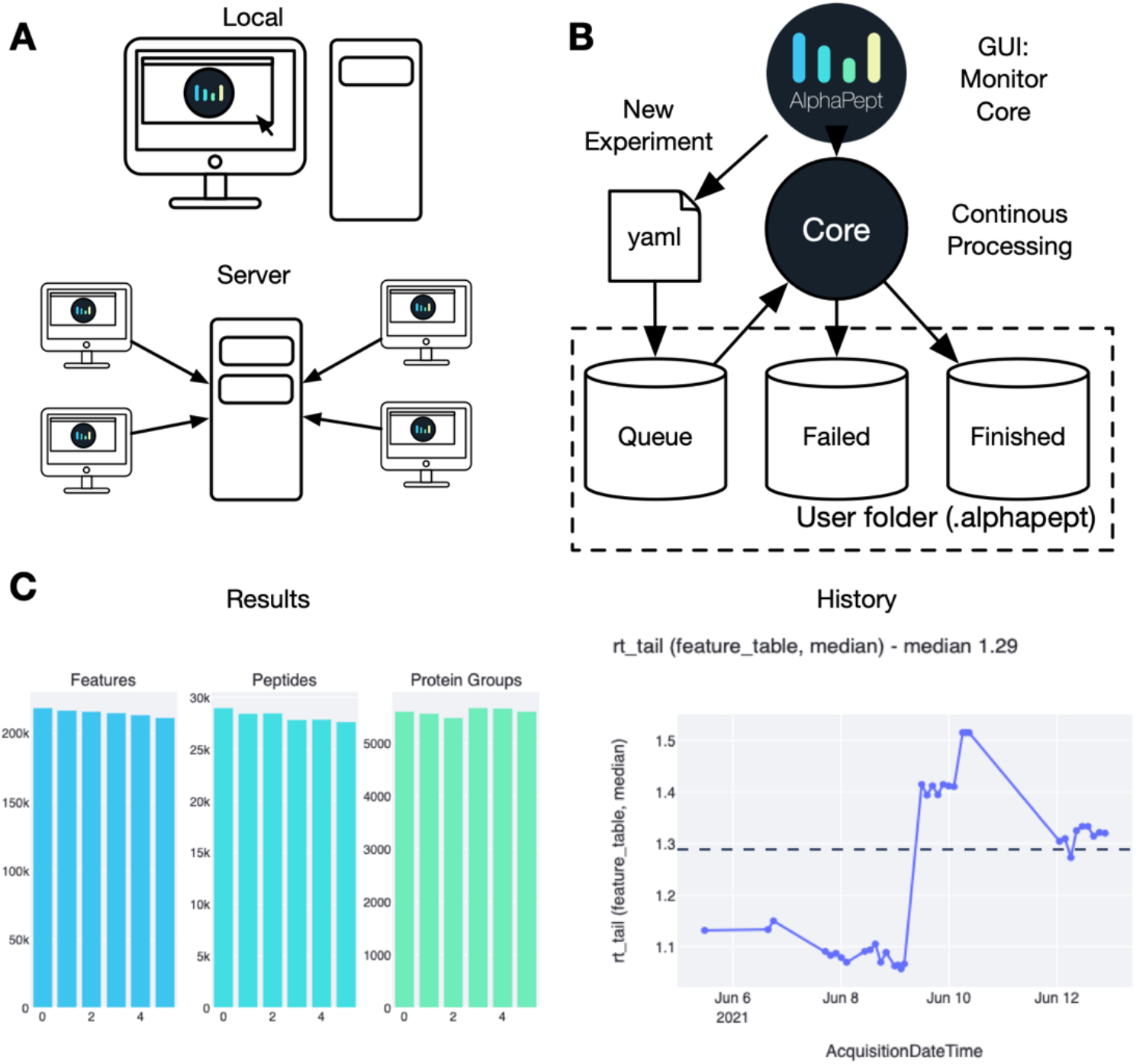
Alphapept user interface, workflow management, deploying and integrating. **A** The AlphaPept GUI is based on a server architecture that can be installed on a workstation and used locally. Additionally, it can be installed on a server and accessed remotely from multiple workstations in the network. **B** AlphePept processing pipeline. The AlphaPept GUI creates three folders for its processing system. New experiments are defined within the interface and saved as YAML files in the Queue folder with automatically triggered processing. **C** Example plots from the History and Results Tab in AlphaPept: Overview of the number of features, peptides and protein groups per injected sample (left panel). Graphing retention time tailing as a function of acquistion date, as an illustration of using AlphaPept for quality assurance.

Adding server functionality typically comes at the cost of maintaining a dedicated API and infrastructure. For AlphaPept we make use of a very recent but already very popular Python package called streamlit (www.streamlit.com), which was developed to facilitate the sharing of machine learning models. By only adding one additional Python package, we have access to a powerful and responsive server infrastructure. Here, the web interface serves merely as an input wrapper to gather the required settings and display results and starts the AlphaPept processing in the background.

### AlphaPept workflow management system

Importantly, the server-based user interface extends the processing functionality of AlphaPept from only processing individual experiments to a continuous processing and monitoring framework. The core processing function of AlphaPept accepts a dictionary-type document to process an experiment, with defined parameters per setting. To store these settings, we chose YAML, a standard human-readable data-serialization language, resulting in files of only a few kilobytes in size. This ensures that they can be modified programmatically and easily checked with common editors.

The settings structure is used by the AlphaPept GUI to build a folder-based workflow management system. It creates three folders in the user folder (‘Queue’, ‘Failed’, and ‘Finished’) and monitors them for new data. When defining a new experiment within the GUI, a settings YAML file is created in the Queue folder, and the core function will start processing. This allows defining multiple experiments, which will then be processed one after another. YAML files of processed runs will be moved to the ‘Finished’ or ‘Failed’ folder (Fig. 8B).

We chose this folder-based processing queue as this allows manual inspection of the processing queue by simply checking the files in the folders. Furthermore, computational alterations of the processing queue are straightforward by writing custom scripts that copy settings files generated elsewhere to the queue folder. AlphaPept has a file watcher module that can monitor folders for new raw files and automatically add them to the processing queue immediately after acquisition is finished. Its modular structure can easily be extended with custom code for integration into larger processing environments with database-based queuing systems. Refer to the interface notebook, which calls the wrapper function and allows customization of the pipeline.

### Visualization of results and continuous processing

For visualization of tabular or summary statistics results, our streamlit application utilizes the ‘Finished’ folder structure where it stores readily accessible summary information of previously processed files (Fig. 8C). AlphaPept has a History tab that compiles these previous results to show performance over time or across analyzed MS runs (Fig. 8D). Here, the user can choose to plot various summary statistics such as identified proteins or peptides as well as chromatographic information such as peak width or peak tailing. As a particular use case, this provides a standard interface which allows instant QC run evaluation in combination with the file watcher.

To inspect an individual experiment, AlphaPept’s browser interface can also plot identification and quantification summary information. Furthermore, basic data analysis functions such as volcano or scatter plots and Principal Component Analysis (PCA) are provided. This is based on streamlit and scikit-learn functionality and can therefore be readily extended. AlphaPept exports the analysis results (quantified proteins and peptides) in tabular format to the specified results path so that it can be readily used for other downstream processing tools such as Perseus (Tyanova et al. 2016) or the recently introduced CKG (Santos et al. 2020).

### AlphaPept deployment and integration

The utility of a computational tool critically depends on how well it can be integrated into existing workflows. To enable maximum flexibility and to address all major use cases, AlphaPept offers multiple ways to install and integrate it.

First, we provide a one-click installer solution that is packaged for a standard Windows system obviating additional installation routines. It provides a straightforward interface to the web-based GUI. We chose Windows for the one-click solution as it is the base OS for the vendor-provided acquisition and analysis software and most users. The one-click installation also has a command-line interface (CLI) for integration into data pipelines.

Next, AlphaPept can be used as a module in the same way as other Python packages. This requires setting up a Python environment to run the tool, which also contains all the functionality of the previously described CLI and GUI. Compared to the Windows one-click installer, the Python module extends the compatibility to other operating systems. While Python code is in principle cross-platform, some third-party packages can be platform bound, such as the Bruker feature finder or DLLs required to read proprietary file types. The modular nature of the AlphaPept file system allows to preprocess files and continue the analysis on a different system (e.g., feature finding and file conversion on a Windows acquisition PC and processing on a Mac system).

Finally, the Python module makes the individual functions available to any Python program. This is particularly useful to integrate only parts of a workflow in a script or to optimize an individual workflow step. Besides the nbdev notebooks that contain the AlphaPept core code, we provide several sandboxing Jupyter Notebooks that show how individual workflow steps can be called and modified. In this way, AlphaPept allows the creation of completely customized workflows.

### AlphaPept processing times

To give the reader an impression of typical processing timings for each of these deployment variants, we ran AlphaPept on various hardware for several use cases: laptop, office PC, workstation and cloud (Table 1). AlphaPept can be readily employed with cloud providers such as Amazon Web Services. We tested our default testing pipeline (see timing table below) on two different Amazon EC2 instances (t3a.2xlarge: 0.42 Eur/h and t3.xlarge: 0.22 Eur/h), an incurred computational costs of 0.22 and 3.82 Euros for one 120 min Orbitrap HeLa file and 8 timsTOF files, respectively, when processed in a European location. Computational costs can be further improved by choosing resource-optimized hardware or buying compute power in advance.

For a typical proteomics laboratory, we envision AlphaPept running in continuous mode, to automatically process all new files. This allows continuous feedback about experiments while drastically speeding up computation when subsequently combining multiple processed files into experiments and experiments into an overall study, because the computational steps that do not change (e.g., raw conversion, database generation or feature finding) can be reused. To illustrate this, the test set with 8 Bruker files from PXD010012 takes 194 minutes on a Workstation with preprocessing and 23 minutes when using preprocessed files.

**Table 2:** Running times of AlphaPept for various hardware (timings in minutes) * IRT = low complexity mixture of peptides (internal retention time standard) ** to process Bruker files on Mac Os X, we preprocessed them on Windows

|  |  | **Laptop**<br>Macbook Pro<br>macOS Big Sur<br>i9 2.3 GHz x 8<br>32 Gb RAM | **Office Pc**<br>Optiplex 7080<br>Windows 10<br>i9 3.7 GHz x10<br>64 Gb RAM | **Workstation**<br>Custom<br>Windows 10<br>i9 3.5 GHz x12<br>128 Gb RAM | **Cloud I**<br>AWS (t3a.2xlarge)<br>Windows Server<br>EPIC 2.2 GHz x4<br>32 Gb RAM | **Cloud II**<br>AWS (t3.xlarge)<br>Windows Server<br>XEON 2.4 GHz x2<br>16 Gb RAM |
| --- | --- | --- | --- | --- | --- | --- |
| IRT Sample*<br>(Thermo) | Full | 1 | 1 | 2 | 3 | 2 |
| HeLa 120 min<br>(Thermo) | Full | 23 | 16 | 19 | 40 | 41 |
|  | Preprocessed | 6 | 4 | 5 | 11 | 12 |
| PXD006109 - 6<br>files (Thermo) | Full | 36 | 17 | 21 | 46 | 73 |
|  | Preprocessed | 30 | 8 | s | 18 | 24 |
| IRT Sample<br>(Bruker) | Full | ** | 1 | 2 | 3 | 2 |
| HeLa 120 min<br>(Bruker) | Full | ** | 57 | 111 | 131 | 399 |
|  | Preprocessed | 6 | 6 | 7 | 16 | 19 |
| PXD010012 - 8<br>files (Bruker) | Full | ** | 242 | 194 | 790 | 893 |
|  | Preprocessed | 62 | 24 | 23 | 85 | 132 |

Being able to import AlphaPept as a Python package also lowers the entry barrier of proteomics analysis workflows for individual researchers and laboratories with little computational infrastructure, as it makes it compatible with platforms like Google Colab, a free cloud-based infrastructure built on top of Jupyter notebooks with GPUs. This allows processing without having to set up software on specialized hardware and allows direct modification of the underlying algorithms. We provide an explanatory notebook for running a workflow on Google Colab, including a 120 min HeLa example file that has been convered on the Windows acquisition computer. This also highlights how the modular HDF5 file format allows us to move the MS data between operating systems.

## DISCUSSION

Here we have introduced AlphaPept, a computational proteomics framework where the relevant algorithms are written in Python itself, rather than Python being used only as a scripting layer on top of compiled code. This architectural choice allows the user to inspect and even modify the code and enables seamless integration with the tools of the increasingly powerful and popular Python scientific ecosystem. The major drawback of such an approach would have been the slow execution speed of pure Python, however extensive use of the Numba just in time compiler – on multiple CPUs or a GPU – makes AlphaPept exceptionally fast, as we have shown in this manuscript. Together with the use of recently developed browser-based deployment, AlphaPept covers the full range of potential users from novice users to systems administrators wishing to build large cloud pipelines.

A related and important design objective of AlphaPept was to enable a diverse user community and invite community participation in its further development. To ensure quality, reproducibility and stability, we implemented a large suite of mechanisms from unit through end-to-end tests via automatic deployment tools. This in turn allows us to streamline the integration of community contributions after rigorous assessment. Furthermore, GitHub provides state-of-the-art tools and mechanisms to allow the effective collaboration of diverse and dispersed developer communities.

Currently, AlphaPept provides functionality for DDA proteomics but we are in the process of enabling analysis of DIA data, ultra-fast access to and visualization of ion mobility data (AlphaTims, https://github.com/MannLabs/alphatims), deep learning for predicted peptide properties and improved quantification, all made possible by its modular design.

One of the large goals of AlphaPept is to ‘democratize’ access to computational proteomics. To this end, besides implementation in Python, we adopted the ‘literate programming’ paradigm which integrates documentation and code. We adopted the nbdev package, providing both beginner and expert computational proteomics researchers with an easy and interactive ‘on ramp’. In our case this takes the form of currently 12 Jupyter notebooks dealing with all the major sub tasks of the entire computational pipeline from database creation, raw data import all the way to the final report of the results. We imagine that students and researchers with novel algorithmic ideas can use this paradigm to add their functionality in a transparent and efficient manner, without having to re-create the entire pipeline. This could especially enable increasingly powerful machine learning and deep learning technologies to be integrated into computational proteomics (Torun et al. 2021; Wen et al. 2020; Meyer 2021).

## Acknowledgements

We thank Sven Brehmer, Wiebke Timm, Konstantin Schwarze and Sebastian Wehner from Bruker Daltonik for providing support with the feature finder for Bruker data. Further, we thank Andreas Brunner, Igor Paron, Patricia Skowronek and Mario Oroshi for providing sample files and descriptions and feedback on the QC pipeline. Xie-Xuan Zhou contributed to discussions and testing. We are grateful of the feedback, testing and support from our group members and colleagues at OmicEra Diagnostics GmbH for testing.

## abbreviations

API: (application programming interface)
CLI: (command-line interface)
DDA: (data-dependent acquisition)
DIA: (data-independent acquisition)
FDR: (false discovery rate)
GPU: (graphical processor unit)
GUI: (graphical user interface)
HDF5: (hierarchical data format 5)
JIT: (just-in-time)
L-BFGS-B: (Broyden–Fletcher–Goldfarb–Shanno)
ML: (machine learning)
MS: (mass spectrometry)
MS/MS: (tandem mass spectrometry)
PASEF: (Parallel Accumulation–Serial Fragmentation)
PRM: (parallel reaction monitoring)
PSM: (peptide spectrum match)
QC: (quality control)
RF: (random forest)
SLSQP: (sequential least squares programming)
trf: (Trust Region Reflective algorithm).

## Software and Data availability

AlphaPept is fully open-source and is freely available under an Apache license at https://github.com/MannLabs/alphapept. All data is available on GitHub or the Max-Planck datashare as test data. Each notebook / file contains respective download links for the files used.The results in this manuscript were obtained with AlphaPept version 0.3.26 if not otherwise indicated.

## Author contributions

MM and MTS conceived the core idea of the AlphaPept framework and MM wrote the first iteration of the search algorithm. MTS wrote the Thermo feature finder, quantification and downstream processing modules, code structure and user interface. EV contributed file importing functionality. IB extended the scoring functionality with ML and FDR control. SW added HDF file handling, revised the general code structure and added performance functions. WFZ and CA contributed and improved quantification. JS critically reviewed testing and documentation. RI and MG contributed to GPU support and code acceleration. All authors contributed ideas, performed testing and wrote the manuscript.

## EXTENDED METHODS

### Notebook availability

All notebooks are available in the repository on GitHub. The documentation created based on the notebooks is available here: https://mannlabs.github.io/alphapept/. Additional information about code not covered in the Notebooks presented here can be found in the Documentation (https://mannlabs.github.io/alphapept/additional_code.html). A cloud hosted Notebook with an example data file is provided at the free Google Colab site: https://colab.research.google.com/drive/163LTlyzBCDgyCkSJiikbmsnny_EiQ7SG?usp=sharing

### MongoDB Dashboard

The continuous integration pipeline has the action “Performance test pyinstaller”. This action freezes the current Python environment into an executable and runs the test files. The results of these tests are uploaded to a noSQL database (MongoDB) for the tested version number. Key performance metrics are visualized in charts here: https://charts.mongodb.com/charts-alphapept-itfxv/public/dashboards/5f671dcf-bcd6-4d90-8494-8c7f724b727b

#### timsTOF and Orbitrap HeLa samples

The test files comprise representative single run analyses of complex proteome samples. Human HeLa cancer cells were lysed in reduction and alkylation buffer with chloroacetamide as previsouly described (Kulak et al. 2014), and proteins were enzymatically digested with LysC and trypsin. The resulting peptides were de-salted and purified on styrenedivinylbenzene reversed-phase sulfonate (SDB-RPS) StageTips before injection into an EASY nLC 1200 nanoflow chromatography system (Thermo Scientific). The samples were loaded on a 50 cm × 75 μm column packed in-house with 1.9 μm C_18_ beads and fitted with a laser-pulled emitter tip. Separation was performed during 120 min with a binary gradient at a flow rate of 300 nL/min. The LC system was coupled online to either a quadrupole Orbitrap (Thermo Scientific Orbitrap Exploris 480) or a trapped ion mobility – quadrupole time-of-flight (Bruker timsTOF Pro 2) mass spectrometer. Data were acquired with standard data-dependent top15 (Orbitrap) and PASEF methods (timsTOF), respectively.

#### timsTOF and Orbitrap iRT samples

11 iRT peptides (https://biognosys.com/product/irt-kit/) were separated *via* a 5.6 min Evosep gradient (200 “samples per day”) yielding test data with low complexity, that facilitated quick testing of computational functionality. An Evosep One liquid chromatography system (Evosep) was coupled online with a trapped ion mobility spectrometry (TIMS) quadrupole time-of-flight (TOF) mass spectrometer (timsTOF pro, Bruker Daltonics). iRT standards (Biognosys) were loaded onto Evotips according to the manufacturers’ instructions and separated with a 4 cm × 150 μm reverse-phase column with 3 μm C_18_-beads (Pepsep). The analytical column was connected with a zero-dead volume emitter (10 μm) placed in a nano-electrospray ion source (CaptiveSpray source, Bruker Daltonics). Mobil phase A contained 0.1 vol% formic acid and water and mobil phase B of 0.1 vol% formic acid and acetonitrile. The sample was acquired with the dda-PASEF acquisition mode. Each topN acquisition mode contained four PASEF MS/MS scans and the accumulation and ramp time were both 100 ms. Only multiply charged precursors over the intensity threshold of 2500 arbitrary units (a.u.) and within a m/z-range of 100 – 1700 were subjected to fragmentation. Peptides that reached the target intensity of 20,000 a.u. were excluded for 0.4 min. The quadrupole isolation width was set to 2 Th below *m/z* of 700 and 3 Th above a *m/z* value of 700. The ion mobility (IM) range was configured to 0.6 – 1.51 Vs cm^−2^ and calibrated with three Agilent ESI-L TuneMix Ions (*m/z*, IM: 622.02, 0.98 Vs cm^−2^; 922.01, 1.19 Vs cm^−2^; 1221.99, 1.38 Vs cm^−2^). The collision energy was decreased as a function of the ion mobility, starting at 1.6 Vs cm^−2^ with 59 eV and ending at 0.6 Vs cm^−2^ with 20 eV.

